# A comprehensive analysis of rheumatoid arthritis B cells reveals the importance of CD11c^+ve^ double-negative-2 B cells as the major synovial plasma cell precursor

**DOI:** 10.1101/2023.02.15.526468

**Authors:** Elinor Wing, Catherine Sutherland, Katherine Miles, David Gray, Carl Goodyear, Thomas Otto, Stefan Breusch, Graeme Cowan, Mohini Gray

**Author notes:** **<u>Author Contributions</u>** MG, EW, KM and GC designed and/or carried out experiments. EW, CS and GC undertook bioinformatics analysis. MG, CG, TO and GC supervised EW. MG conceived of project, obtained grant funding, and with EW, wrote the manuscript.

## Abstract

B cells are key pathogenic drivers of chronic inflammation in rheumatoid arthritis (RA). There is limited understanding of the relationship between synovial B cell subsets and pathogenic antibody secreting cells (ASCs). This knowledge is crucial for the development of targeted therapies. Here, we combine flow cytometry of circulating B cells with single-cell RNA and paired repertoire sequencing of over 27,000 synovial B cells from patients with established RA. Twelve B cell clusters were identified including previously recognised subsets, and a novel cluster that strongly expressed heat shock proteins. All lineages identified by trajectory analysis contribute to the DN2 B cell population, which is the major precursor to synovial ASCs. This was further supported by B cell receptor (BCR) lineage analysis, which revealed clonal relationships between DN2 cells and ASCs. This study advances our understanding of B cells in RA and reveals the origin of pathogenic ASCs in the RA synovium.

## Introduction

Rheumatoid arthritis (RA) is the commonest autoimmune mediated inflammatory arthritis affecting about 1% of the world’s population. As well as chronic, severe pain and early joint destruction, patients experience marked fatigue, accelerated cardiovascular disease and premature death (1, 2). Despite the heterogenous clinical presentation, the importance of B cells in disease pathogenesis has been established, and further reinforced by the success of B cell depleting therapies (BCDT) (3, 4). Pathogenic B cells contribute to the chronic synovial inflammation in at least three ways; cytokine secretion, antigen presentation, and autoantibody production (5). In RA, B cell dysregulation occurs long before clinical disease onset, and BCDT has been shown to delay the onset of clinical disease in preclinical patients (6, 7). Circulating autoantibodies to self-antigens that include, but are not limited to, citrullinated auto-antigens (ACPA) are present years before arthritis becomes clinically evident (8, 9). Immune complexes formed with self-antigens become trapped within the cartilage of synovial joints, activating innate immune cells, that drive inflammation and joint damage. Within the diseased synovium the most common histological pathotype is dominated by the presence of B cells, that both serve as the main antigen presenting cell (APC) to T cells and secrete pro-inflammatory cytokines (10).

Although BCDT can be beneficial in treating RA, it is non-specific, depleting all CD20^+ve^ B cells. This results in the loss of not only pathogenic B cells, but also protective B cells, which are vital for tissue homeostasis and infection control. The importance of the latter was borne out during the SARS-CoV-2 pandemic where mortality and morbidity was significantly increased in patients with autoimmunity, who had received BCDT prior to infection. In addition, the protective effect from SARS-CoV-2 vaccination was reduced in these patients (11-14). This highlights the need for more selective therapies that target only pathogenic B cells. Such therapies can only be developed when the identity of autoimmune B cells, particularly those that serve as precursors to synovial CD20^-ve^ antibody secreting cells (ASCs) is known and so delineating these cells is an urgent priority.

Single cell analysis of the rheumatoid synovium has previously identified 4 B cell subsets, including naïve B cells, memory, double negative B cells and terminally differentiated antibody secreting cells (ASCs) (15). However, the inter-relatedness of these subsets is poorly understood and importantly, the major precursor to rheumatoid synovial CD20^-ve^ ASCs has not been established.

“Double negative” (DN) refers to a subset of antigen experienced B cells, that lack expression of both CD27 and IgD. They are further divided into DN1 and DN2 cells of which the latter have high expression of CD19, CD11c and the transcription factor T-bet, but lack the expression of CD21 and CXCR5. Whilst DN1 B cells represent the majority of DN B cells in healthy subjects, DN2 B cells are increased in the elderly, during chronic infection (including hepatitis C, malaria, and HIV) (16-21) and following influenza vaccination (22, 23). Most recently they have been linked to severe and fatal outcomes from COVID-19 infection, suggesting that they have prominent pro-inflammatory functions (24).

DN2 B cells substantially increased in the circulation of patients with active systemic lupus erythematosus (SLE), where they are extra-follicular precursors to antibody secreting cells (ASCs) and correlate with increased disease activity and poorer outcomes (21). They have also been reported in patients with Sjogren’s syndrome, multiple sclerosis, and Crohn’s disease (25-28). We previously observed a significant increase in the frequency of circulating class switched IgG^+ve^ DN2 cells in RA patients. These B cells also expressed significantly fewer mutations in the B cell receptor (BCR) and were enriched in the synovium, suggesting an extrafollicular origin (29). We considered that naïve B cells entering the synovium were induced to mature into antibody secreting cells via the generation of DN2 cells in extrafollicular areas. The aim of this study was to provide a detailed assessment of circulating and synovial B cells in rheumatoid arthritis and to identify the main precursors to pathogenic ASCs in the diseased synovium. In addition, by utilising single cell sequencing and RNA velocity we aimed to define the interrelatedness of the identified synovial B cell subsets.

## Results

### DN2 cells are enriched in the blood and synovium of RA patients

To fully characterize rheumatoid B cells, a cohort of 29 RA patients and 15 matched healthy controls were selected for full spectrum flow cytometry (Supplementary Table 1). We studied all known B cell subsets in the blood (Supplementary Table 4). In agreement with our previous studies, double negative-2 (DN2) cells were increased in the circulation of RA patients when compared to healthy controls (means with 95% confidence intervals were 1.51% ± 0.3 vs 0.76% ± 0.38, P=0.0003) (Fig. 1A-B). In addition, DN2 B cell precursors, called activated naïve B cells (Anav), that are IgD^+ve^ CD27^-ve^ CD21^-ve^ CD24^-ve^ CD38^-ve^ (21), were almost doubled in frequency in the blood of RA patients compared to the healthy controls (2.35% ± 0.76 vs. 1.13% ± 0.45, P=0.0091) (Fig. 1C). The characteristic surface markers of DN2 cells, that delineate them from both naïve and memory B cells, including CD11c, CD21, CD24 and CD73, established their full phenotype and the expression of both HLA-DR and CD86 suggested an enhanced capacity to act as APCs (Fig. 1D). Importantly, DN2 B cells constituted a significantly higher proportion of synovial B cells compared with that of DN2s in the circulating B cell population. (3.02% ± 1.97 vs. 12.0% ± 6.46, P=0.0257) (Fig. 1E). In health, the frequency of circulating DN1 B cells predominates over DN2 B cells. However, in RA patients the ratio of circulating DN2 to DN1 B cells was increased compared to healthy controls (0.338 ± 0.106 vs. 0.101 ± 0.04, P<0.0001), due to the expansion of the DN2 B cell population (Fig. 1F). The proportion of DN2 cells in the circulation did not correlate with age, suggesting that these cells are distinct from the more broadly described age-associated B cells (Fig. 1G).

**Figure 1:**
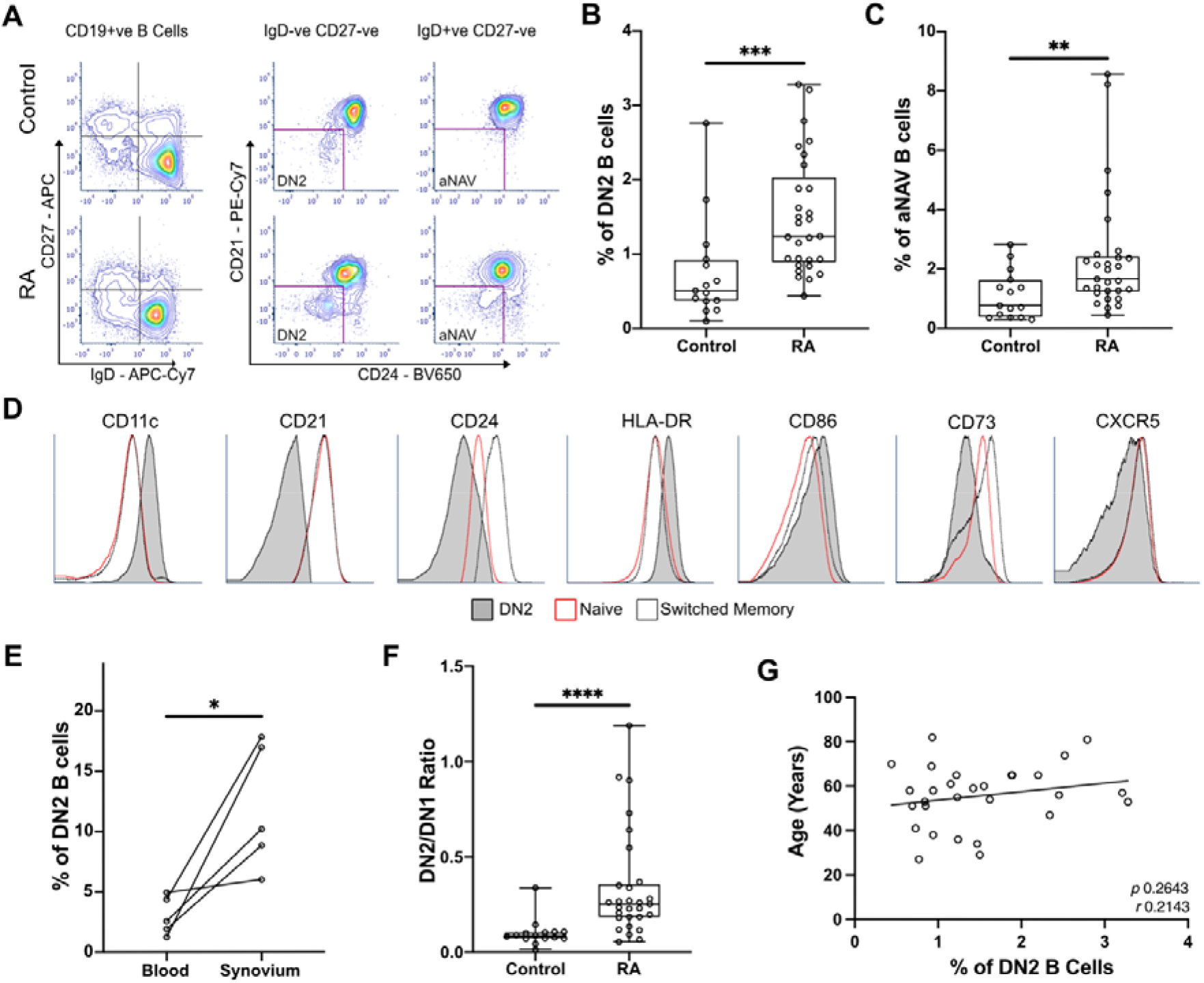
DN2 cells are significantly enriched in the blood and synovium of RA patients. A) Peripheral blood mononuclear cells (PBMCs) were stained for flow cytometry, live CD19+ve B cells were gated before selecting either the double negative (DN; CD27^-ve^ IgD^-ve^) or the naïve (CD27^-ve^ IgD^+ve^) cells. The expression of CD21, CD24, and CD38 (not shown) were used to gate to double negative 2 (DN2) and activated naïve B cells (aNAV). Representative plots show the expansion of DN2 and activated naïve B cells (aNAV) in RA. B) Percentage of DN2 cells within the CD19^+ve^ B cell population. n=29 RA patients and 15 healthy controls. P=0.0003 by Mann-Whitney test. C) Percentage of aNAV cells within the CD19^+ve^ B cell population. n=29 RA patients and 15 healthy controls. P=0.0091 by Mann-Whitney test. D) Representative histogram plots show the expression of relevant surface markers on DN2 (filled grey), resting naïve (red line), and switched memory B cells (dotted line). E) Flow cytometry of paired peripheral blood and synovium. DN2 cells are significantly increased in the synovium of RA patients compared to blood (n=5). P=0.0257 by Paired t test. F) The ratio of DN2:DN1 B cells is significantly increased as a result of DN2 B cell expansion. P<0.0001 by Mann-Whitney test. G) Scatterplot of the percentage of DN2 cells and age of RA donors. Spearman’s r coefficient. * P ≤ 0.05, ** P ≤ 0.01, and *** P ≤ 0.001.

### Single-cell sequencing of synovial B cells reveals the heterogeneity of B cells in the rheumatoid joint

Highly purified synovial B cells (CD19^+ve^ CD14^-ve^ CD3^-ve^) were isolated from the joints of 3 seropositive patients with established RA at arthroplasty (Supplementary Fig. S1A). Single cell RNA sequencing (scRNA-seq) was carried out using the 10X Genomics Chromium platform to obtain paired transcriptomic and BCR sequence data for each cell (Fig. 2A). In total, data for 27,053 synovial B cells were analysed, 7,825 of which had productive heavy and light chain BCR sequence information. These cells clustered into 12 distinct subsets that were annotated according to known markers and signatures (Fig.2B, Supplementary Fig. S1B-C).

**Figure 2:**
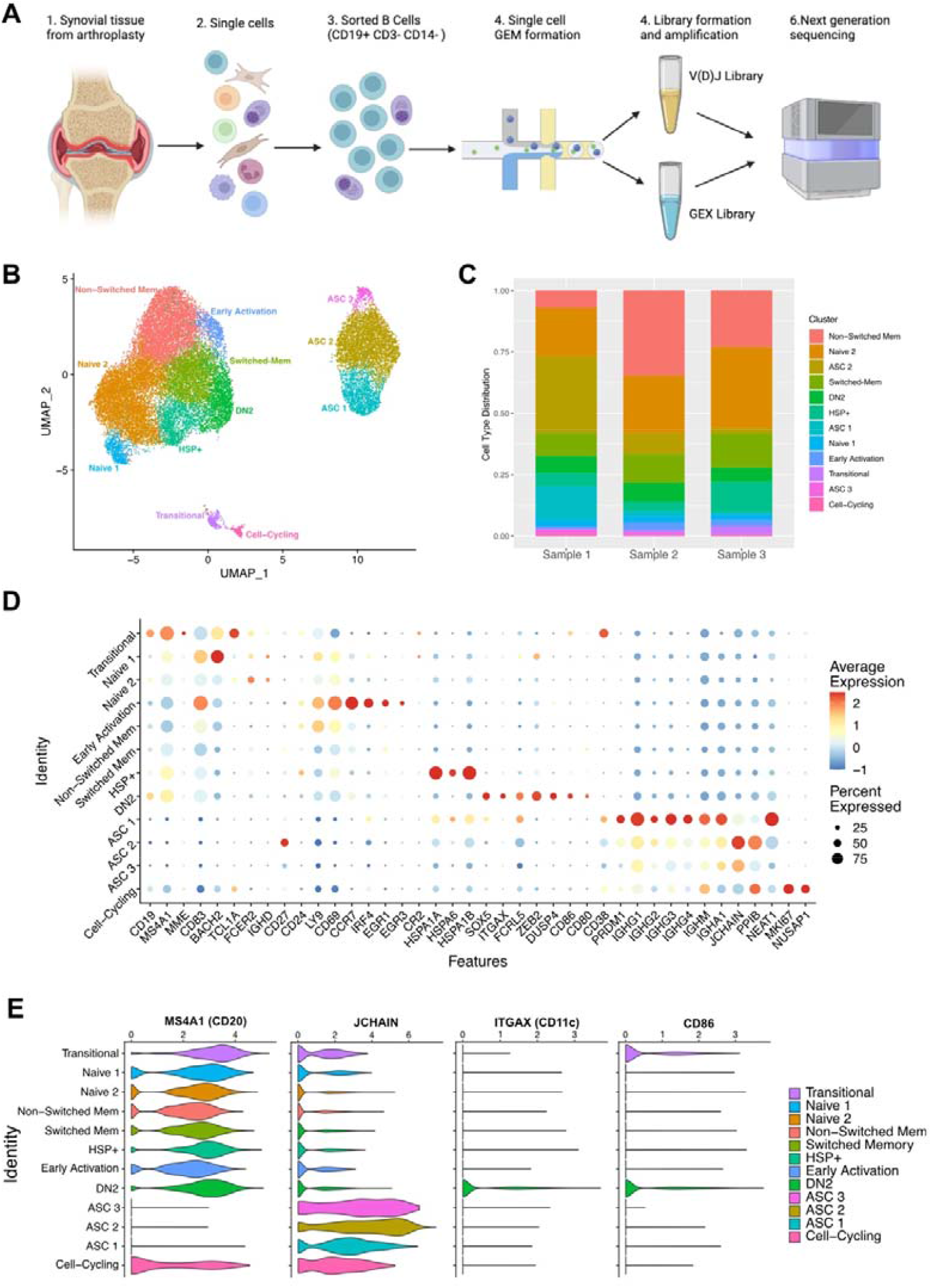
Single-cell sequencing of synovial B cells unveils the heterogeneity of B cells in the rheumatoid joint. A) Overview of sample preparation: CD19+ve B cells were isolated from the synovial tissue of RA patients (n=3) before single cell RNA-sequencing with paired BCR sequencing. B) Uniform manifold approximation and projection (UMAP) projection of all 27,053 synovial B cells from RA patients (n=3). Unsupervised clustering identified 12 clusters, the B cell subsets were annotated using the expression of known subset markers and gene signatures. DN2, double negative 2; HSP, heat shock protein; ASC, antibody secreting cell. C) Stacked bar plot showing the distribution of the 12 clusters over the 3 samples. D) Dotplot demonstrating the expression of key gene markers for each cluster. E) Violin plots demonstrating the expression of *MS4A1* (CD20), *JCHAIN, ITGAX* (CD11c), and *CD86* across the 12 clusters.

Naïve B cells (split into Naïve 1 and Naïve 2) comprised just over a quarter of all B cells (7,295 B cells; 26.9% of total) (Fig. 2B and 2C). *IGHD* expression was found primarily in these clusters alongside minimal expression of *CD27*, matching the canonical surface markers used to identify naïve B cells by flow cytometry. Compared to the Naïve 2 cluster, the Naïve 1 cluster differs by the increased expression of *BACH2* and *CD83* and the reduced expression of *FCER2* (CD23) (Fig. 2D), suggesting that the Naïve 1 B cells may be recently activated.

Memory B cells were identified by the increased expression of *CD27* and totaled 9,251 (34.2%). This population was further split into switched or unswitched according to BCR isotype. A small population of B cells (604 cells, 2.2%) expresses high levels of *CCR7* compared to the other clusters and were defined as being in a state of early activation. The increased expression of *CD83, IRF4, CD69, EGR1* and *EGR3* also suggests that cells within this cluster have been recently activated (Fig. 2D).

6.9% (1,891 of all B cells) expressed high levels of *ITGAX* (CD11c), as well as *FCRL5, CD86, SOX5*, and *MS4A1* (CD20) (Fig. 2D & 2E), which are hallmarks of DN2 cells. Cells within this cluster lack expression of *CD27, FCER2* (CD23) and *CR2* (CD21), which is consistent with the DN2 population identified using full-spectrum flow cytometry. *JCHAIN*^+ve^ ASCs made up just over 21% of the total B cell population (5,786 cells) and encompassed three separate clusters (ASC 1, ASC 2, and ASC 3). As expected, the ASC clusters sit separately from the other B cell clusters, expressing high levels of *JCHAIN* and *PRDM1* (BLIMP1). The latter is an important regulator of ASC differentiation, that promotes the expression of *XBP1* and represses B cell transcription factors. Three separate ASC clusters were identified and showed distinct levels of *CD38, CD27, PRDM1* and *NEAT1* expression (Fig. 2D). The higher expression of *NEAT1*, a MYC-regulated long noncoding RNA (lncRNA) that promotes proliferation, in the ASC 1 cluster suggests that these cells are proliferating and have a more plasmablast-like phenotype. The ASC 2 cluster expresses the most *PPIB* (that forms complexes facilitating protein folding) and *JCHAIN*, both of which are involved in the unfolded protein response, that is activated during periods of high antibody production due to the immense stress on the cells. B cells from the ASC 3 cluster have the lowest average expression of ASC related genes implying terminal differentiation and exhaustion (Fig. 2D).

A novel cluster of 1,708 (6.3%) B cells expressed high levels of heat shock protein (HSP) genes and are called henceforth HSP^+ve^ B cells. Although an increase in HSPs has been reported in the synovial fluid of RA patients, the subset of immune cells producing them has not been identified (30). The B cells in the HSP^+ve^ cluster expressed high levels of HSP genes including *HSPA1A, HSPA6, HSPA1B*, and *HSPB1* (Fig. 2D and Supplementary Fig. S3A). The high expression of HSPs is linked to the misfolded protein response and de novo protein folding pathways (Supplementary Fig. S3B).

Smaller additional populations include 361 (1.3%) *MME*^+ve^ (CD10) transitional cells and 157 (0.6%) cell-cycling B cells (Fig.2B). All clusters were present in each of the samples; however, the ASC clusters were primarily composed of cells from Sample 1 (68% of ASCs). The proportion of DN2 cells was similar across the three samples, with RA1 having 6.8%, RA2 having 7.8%, and RA3 having 5.6%. (Fig.2C).

### DN2 cells are a heterogenous group of cells, primed to present antigen and become ASCs

Differential expression analysis showed that the DN2 cluster was enriched in the expression of *ITGAX* (CD11c), *SOX5, ZEB2, DUSP4, FCRL5, LRMP, HOPX, CD86* and *NEAT1*, matching previous reports of DN2 cells in humans and CD21^lo^ B cells in mice (21, 31) (Fig.3A). *SOX5* is a transcription factor linked to the late stages of B cell differentiation and has previously been found to be enriched in CD21^lo^ and atypical memory B cells which share features with DN2 cells (32). *ZEB2* has also been previously linked to autoimmune B cells as it directly binds the promotor of *ITGAX*. We compared this data to FACS sorted bulk RNA-seq data from DN2 cells of SLE patients (21) as well as the homologous mouse CD21^lo^ CD23^lo^ B cell population (31). Projecting the location of cells with a module score in the top 10% onto the UMAP confirmed that these expression patterns overlap with the DN2 cluster in this dataset, as well as the ASC2 and ASC3 clusters (Fig.3B). Further GSEA analysis using the GO Biological pathways revealed strong enrichment for pathways related to antigen processing and presentation, particularly exogenous antigens via MHC Class II, due to the increased expression of CD86 and HLA genes including *HLA-DQA2, HLA-DQB1*, and *HLA-DRB1* (Fig. 3C).

**Figure 3:**
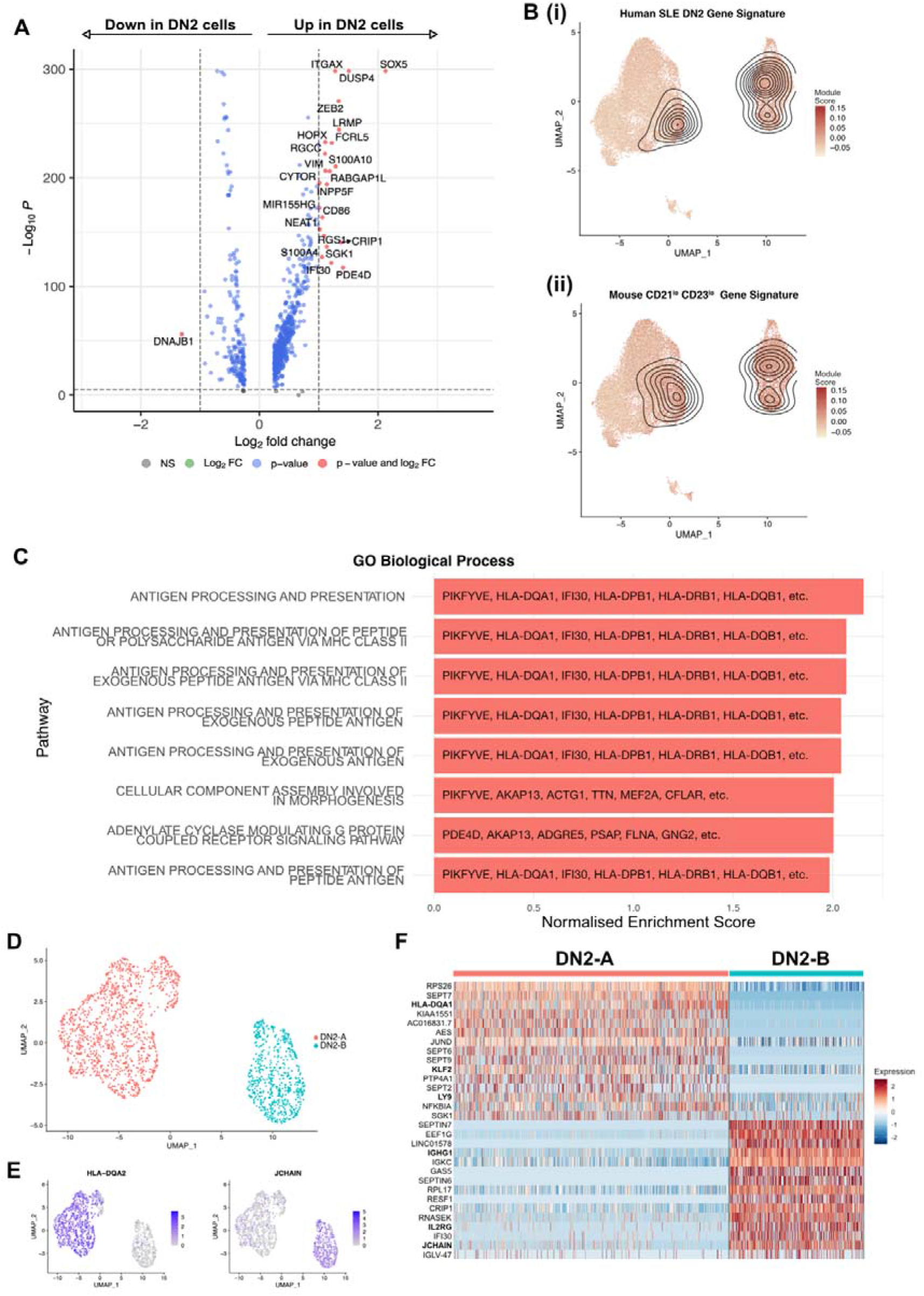
DN2 cells are a heterogenous group of cells, primed to present antigen and become ASC. A) Volcano plot showing the transcriptome analysis DEGs (red, log2FC is >|1| and P value > 10e-6; blue, log2FC is <|1| and P value > 10e-6; grey, nonsignificant genes) of DN2 cells compared to all other non-ASC cells. B) UMAP projection of scRNA-seq data from 27,053 synovial B cells from RA patients (n=3) with the module scores for (i) human SLE DN2 cells and (ii) CD21^lo^ CD23^lo^ B cells from mice, the contour overlay represents the position of the top 10% of module scores. C) The top 8 pathways enriched in the DN2 cells as identified by gene set enrichment analysis using gene sets derived from the GO Biological Process ontology. D) UMAP plot of 1,891 DN2 cells from RA synovial tissue (n=3), unsupervised clustering revealed two distinct populations, DN2-A and DN2-B. E) Expression of HLA-DQA2 and JCHAIN in DN2-A and DN2-B. F) Heatmap of the top 20 up- and downregulated genes between the DN2-A and DN2-B subsets with important genes highlighted.

Re-clustering of the 1,891 DN2 cells further revealed two distinct populations, DN2-A and DN2-B (Fig. 3D). DN2-A differentially expressed *HLA-DQA1*, which affects the loading of peptide antigens onto class II molecules for presentation to T cells, implying that DN2-A B cells are primed to present antigen and activate CD4^+ve^ T cells. DN2-A B cells also expressed higher levels of genes related to cell survival and quiescence such as *KLF2* and *JUND*. In contrast, DN2-B cells expressed higher levels of *JCHAIN, IGHG1, IGHA1*, and *IL2RG* indicating that they are transitioning into ASCs (Fig. 3E-F).

### Trajectory inference shows DN2 cells are ASC precursors in the RA synovium

As the source of autoreactive antibodies that form immune complexes, ASCs are a major contributor to inflammation and tissue damage within the joint. To delineate the developmental processes within the joint and pinpoint the main precursors of ASCs we have used a combination of RNA velocity and trajectory inference. RNA velocity analysis with scVelo (33) revealed that mature B cell clusters evolve into DN2 cells with a subsequent onward flow towards ASC clusters (Fig. 4A). Trajectory inference with Slingshot (34) revealed three separate lineages (Fig. 4B). Lineage 1 starting in the Naïve 1 cluster, transitioning through the non-switched B cell clusters into the switched memory clusters before merging into DN2 cells, and finally differentiating into ASCs. Lineage 2 begins in the Cell-Cycling cluster, to Switched-Memory, to DN2, to ASC. Finally, Lineage 3 starts in the Early Activation cluster before joining Lineage 1 in the Non-Switched Memory cluster to DN2 cells and then to ASCs.

**Figure 4:**
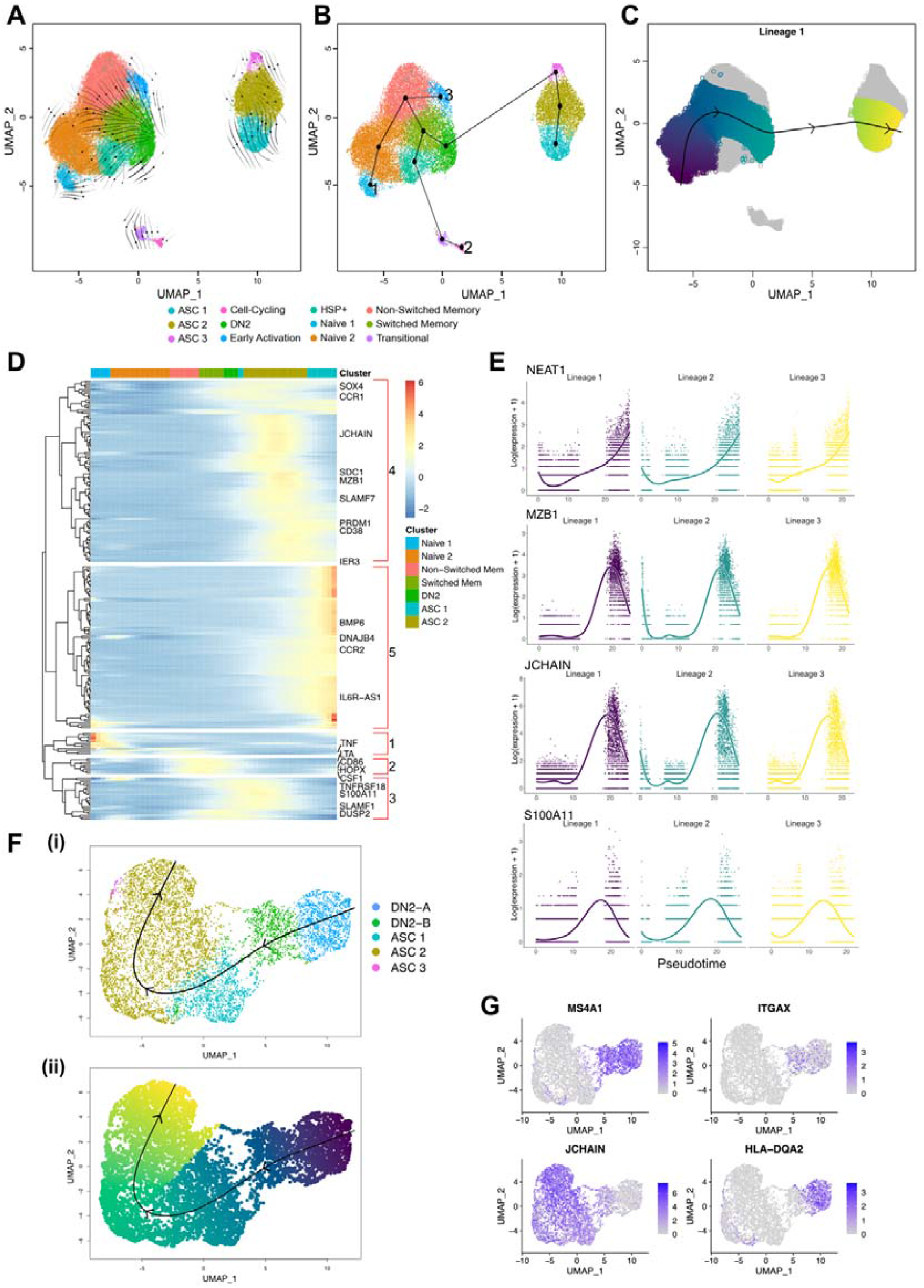
Trajectory inference shows DN2 cells are ASC precursors in the RA synovium. A) RNA velocity stream plot, projected onto the UMAP of scRNA-seq data from 27,053 synovial B cells from RA patients (n=3). B) Minimum spanning tree demonstrating the full lineage structure identified by Slingshot. Lineage 1 starts in the Naïve 1 cluster, Lineage 2 in the Cell-cycling cluster, and Lineage 3 in the Early Activation cluster. C) Lineage 1 identified by Slingshot with pseudotime colouring. D) Top 250 differentially expressed genes in Lineage 1, split into 5 groups. E) Four of the most significant genes identified by comparing the mean expression at the start of the lineage to the end of the lineage. F) Slingshot trajectory identified after clustering 1,891 DN2 cells and 5,786 ASCs from RA synovial tissue (n=3). (i) The trajectory starts in the DN2-A cluster, then becomes DN2-B before transitioning into ASCs. (ii) The lineage identified by Slingshot with pseudotime colouring. G) Expression of *MS4A1* (CD20), *ITGAX* (CD11c), *JCHAIN*, and *HLA-DQA2* in DN2 cells and ASCs.

Changes of gene expression along Lineage 1 can be clustered into 5 groups. The genes in Group 1 peak in expression early in the trajectory in the Naïve 1 cluster. Then, group 2 contains genes that are expressed in mature B cells and includes the cytokine genes *CSF1* and *LTA*. Group 3 contains genes that begin to be upregulated in the DN2 cluster and into the ASCs. This group contains *SLAMF1* that is upregulated upon activation in B cells and promotes proliferation and differentiation into ASCs. Group 4 contains typical ASC gene markers such as *JCHAIN, SDC1* (CD138), *SLAMF7, PRDM1* (BLIMP-1), and CD38 which all peak in expression as the cells begin to transition into ASCs.

Finally, group 5 contains genes which peak at the very end of the differentiation into ASCs, including *DNAJB4* which is involved in ensuring correct protein folding and *BMP6*, that helps to inhibit proliferation in terminally differentiated cells (Fig. 4D).

The expression of several of the important genes for the differentiation of ASCs like *NEAT1, MZB1, JCHAIN* begin to increase in the DN2 cluster (Fig. 4E). *S100A11* increases in the DN2 cluster as they become ASCs (Fig. 4E) and is known to be elevated in the synovial fluid and tissue of RA patients, though its cellular source was previously unknown (35).

To further understand the transition between DN2 and ASC clusters, we carried out clustering of only the DN2 cells (1,891 cells) and the ASCs (5,786 cells) together. Within this subset, Slingshot identified 1 lineage starting at the *HLA-DQA2*^high^ DN2-A cluster, before passing through the *JCHAIN*^high^ DN2-B cluster into the ASC cluster (Fig 4F). When looking at the expression of *MS4A1* and *JCHAIN* there is a clear gradient showing the decrease in CD20 and the increase of *JCHAIN* as the cells differentiate into ASCs (Fig. 4G).

### BCR sequences from synovial B cells show hallmarks of autoimmunity

To further understand the diversity of BCRs present in the inflamed rheumatoid joint, the paired BCR repertoire sequences were evaluated, specifically looking at characteristics linked to autoimmunity. The mean IGH CDR3 lengths were similar across the three samples with Sample 1 having a mean CDR3 length of 18.3 amino acids, Sample 2 18.0, and Sample 3 17.5 amino acids (Fig. 5A). Over a quarter (26.2%) of all IGK CDR3s are over 11 amino acids long, which has previously been linked to autoimmunity and poly-reactivity (36).

**Figure 5:**
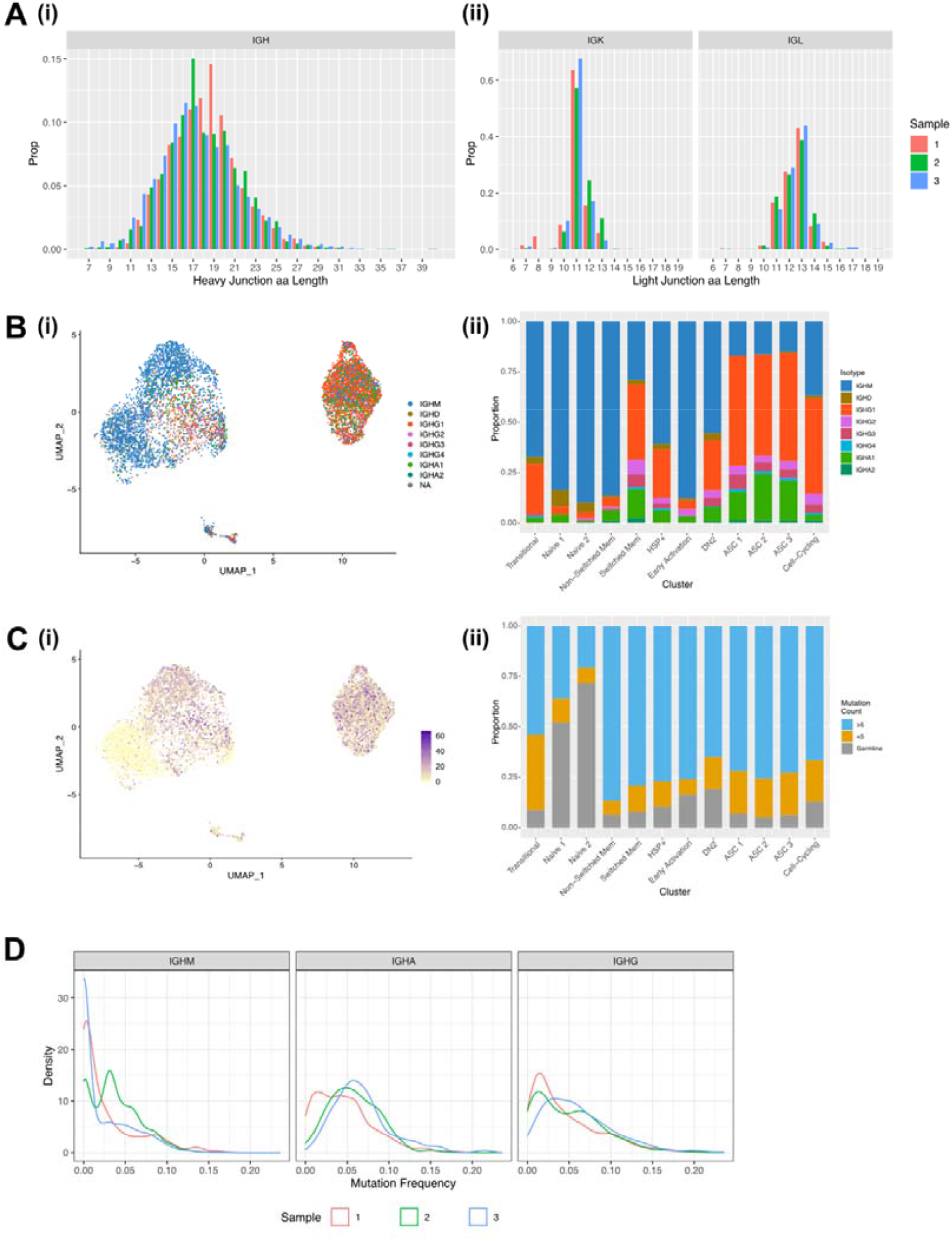
BCR sequences from synovial B cells show hallmarks of autoimmunity. A) CDR3 length distribution for the (i) heavy chain and the (ii) light chains for Sample 1 (red column), Sample 2 (green column), and Sample 3 (blue column). B) (i) UMAP plot of the 7,825 B cells with a productive heavy and light chain coloured by the BCR isotype. (ii) Stacked bar plot showing the distribution of the isotypes across the 12 clusters. C) (i) UMAP plot of the 7,825 B cells with a productive heavy and light chain coloured by the BCR mutation count. (ii) Stacked bar plot showing the distribution of mutation counts across the 12 clusters. D) Distribution of mutation frequencies for IGHM, IGHA, and IGHG for the three RA samples.

As expected, the naïve clusters primarily expressed unmutated IGHM and IGHD with IGHA and IGHG being expressed in the memory and ASC clusters where class switching has occurred after activation. (Fig. 5B). As B cells become activated both the mutation rate and proportion of class-switched cells increases, with this being evident at the border between the Naïve 2 and the memory clusters (Fig. 5B and 5C). Despite the expected increase in mutation rate, unmutated sequences were present in all the clusters. In particular, 19% of the DN2 cells expressed BCR sequences which are identical to germline sequences (Fig. 5C). Low mutation rates persisted in some of the terminally differentiated ASCs, suggesting development out with germinal centres.

For the class-switched sequences there is predominantly a unimodal distribution of mutations peaking around 5% mutation frequency but extending up to 25%. The majority of IGHM sequences remain unmutated or poorly mutated with an average mutation frequency of 3% compared to 5% in IGHG and IGHA. The sharp peak in IGHM Sample 2 comes from a large clonal expansion of an IGHM BCR sequence (Fig. 5D).

### BCR sequences from DN2 cells share identical CDR3 sequences with ASCs

We analysed the BCR sequences in the synovium to investigate clonal relationships. The synovial B cells showed a high degree of clonal expansion, with 62.4% and 40.2% of clones being expanded (clone size ≥[]2) in Samples 1 and 2, respectively. Sample 3 was less clonal, with only 9.1% of clones consisting of at least two cells (Fig. 6A and 6B). To further compare the degree of clonal expansion between the three patients we used the Gini coefficient, a measure of inequality. A Gini coefficient of 0 indicates maximal diversity or an equal abundance of each sequence, whereas 1 indicates extreme inequality. In concordance with the proportion of expanded clones, Sample 1 had the highest Gini coefficient (0.33) followed by Sample 2 (0.21), with Sample 3 having a much lower value (0.03).

**Figure 6:**
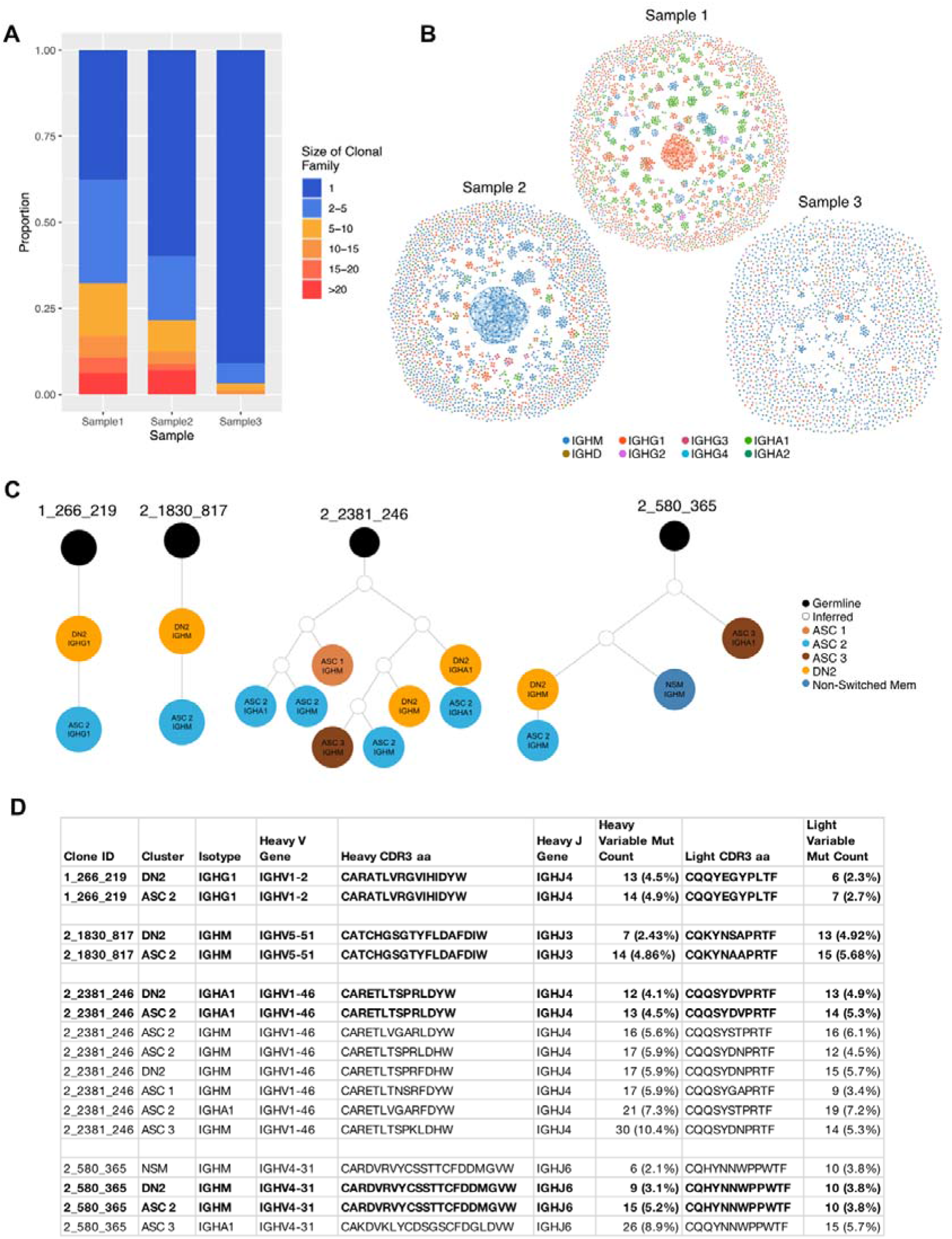
BCR sequences from DN2 cells share identical CDR3 sequences with ASCs. A) Stacked barplot showing the clonal family sizes across the three samples. B) Network graphs of a representative repertoire from each RA patient. Each node represents a cell, coloured according to the isotype, with up to three edges representing the most closely related heavy CDR3 amino acid sequences according to Levenshtein distance. C) Four clonal lineage trees where a DN2 B cell shares an identical heavy CDR3 amino acid sequence with an ASC. Orange circles indicate a cell from the DN2 cluster, dark blue non-switched memory, dark brown ASC 1, bright blue ASC 2, pale brown ASC 3, white inferred sequence and black is germline. D) Characteristics of the 4 clonal families where a DN2 B cell shares an identical junction sequence with and ASC.

There was no relationship between the size of the clonal group and the rate of mutation. The largest clonal group in Sample 1 contained 87 class switched IgG1 cells and the most mutated sequence had just 4 mutations in the heavy variable sequence, whereas a smaller group with 14 cells had up to 52 mutations (Supplementary Fig. S4A).

Having identified (through trajectory inference and RNA velocity) that DN2 cells are a precursor to ASCs, we explored the relationship between the BCR sequences in these clusters. 4 clonal groups had an identical heavy CDR3 amino acid sequence between a DN2 B cell and an ASC (Fig. 6C and 6D). These direct links, ending at an ASC, originated from 4 DN2 cells, 2 Switched Memory cells, 1 Naïve 2 cell and 4 Cell-Cycling cells, demonstrating again that DN2 cells are the primary precursor to ASCs (Supplementary Fig. S4B).

## Discussion

This is the most detailed exploration to date of both circulating and synovial B cells subsets in patients with RA. By combining trajectory inference and RNA velocity we have mapped the maturation of naïve B cells to ASCs. We show that the B cell clusters are intimately linked from naïve through memory to DN2 cells and critically, that DN2 cells serve as the major precursors to synovial ASCs. In addition, DN2 B cells expressed potent APC capacity and are therefore likely to represent an important pathogenic B cell subset that should be targeted for future therapies.

DN2 B cells play an important role in the pathogenesis of a related autoimmune rheumatic disease, SLE, where they are enriched in blood, being epigenetically poised to respond to TLR7 ligands and serve as the pathogenic precursors to ASCs (21). We have previously observed that DN2 B cells were enriched in both the blood and synovium of RA patients (28), and confirmed this finding in a second, independent cohort of 29 patients. Indeed, we found both circulating DN2 B cells and their precursors aNAVs are almost twice as frequent in RA patients. This is supported by previous reports that see an elevation of DN B cells in the blood (37, 38). This increase can be explained by a change in the dominant subtype in RA. We have found that the ratio of DN1 to DN2 B cells is reversed in RA so that the frequency of DN2 B cells as a percentage of all B cells is higher, whilst the DN1 frequency itself does not change. Hence to identify a significant elevation in pathogenic DN2 B cells, all the available markers to differentiate DN1 and DN2 B cells must be included and DN B cells should not be treated as a single population.

DN2 cells were present in the blood of healthy controls, albeit at very low numbers. A recent single cell study assessed the immune compartments of 16 tissues (not including the synovium) from 12 adult donors. They identified a B cell subset referred to as Age Associated B cells (ABCs), that has a similar transcriptional profile to our DN2 cluster, including high expression of *ITGAX* and low expression of *CR2* (CD21) and *CD27* (Supplementary Fig. S2A). They found that DN2 cells were primarily found in the spleen, liver, and bone marrow (32). DN2 cells are clearly important in health but dramatically increase in autoimmune conditions and viral infections, as we have shown in RA and others have demonstrated in patients with SLE, HIV, and COVID-19 (18, 21, 24).

By assessing 27,000 synovial B cells we were able to identify 12 B cell clusters, comprised of 2 naïve, 2 memory, 3 ASC, the DN2 clusters along with 4 much smaller clusters. One of these smaller clusters was entirely novel and expressed heat shock proteins (HSPs) from the HSP70 family. In RA, serum levels of HSP-70 are about twice the level of healthy controls and significantly elevated in the synovium (39). Whilst their main role is to reverse or inhibit denaturation or unfolding of cellular proteins in response to stress, they are associated with an increased frequency of Th17 T cells and the Th17/Treg ratio (40). It’s possible that this B cell cluster may be a significant source of serum and synovial tissue HSPs, given that B cell lines readily release HSP70 via exosomes (41). The role this HSP^+ve^ B cell subset is playing in the pathogenesis of RA is unknown but deserves closer scrutiny in follow up studies.

Two other studies utilising single cell sequencing of rheumatoid synovial B cells have been reported. In the first just over 1000 B cells were available for analysis (from 21 subjects with established RA). Here four synovial B cell subsets were identified, that included naïve, memory, plasmablasts and DN2 B cells, (that were referred to as autoimmune B cells or ABCs in the study) (15). Of note the ABCs were derived largely from tissue samples that were leukocyte rich and expressed the interferon stimulated genes *GBP1* and *ISG15*. As we also selected samples that were leukocyte rich (by flow cytometry), the presence of DN2 B cells may be a particular feature of this synovial pathotype and cannot necessarily be extrapolated to other histological synovial phenotypes, in particular the leukocyte poor and macrophage rich subtypes (10, 42, 43). Of note, leukocyte rich synovium is associated with the highest inflammation scores clinically and so patients with this phenotype are most likely to benefit from targeted B cell depletion (10).

A further study of newly diagnosed, untreated RA patients assessed just under 2000 B cells from 3 RA patients, 2 of whom were seronegative for antibodies directed at citrullinated autoantigens (i.e., ACPA^-ve^). Here there was no enrichment of *ITGAX*+ve (encoding CD11c) or CD27^-ve^ IgD^-ve^ B cells (44). The difference between these observations and our study may relate to the early stage of RA and the limited number of B cells assessed with the predominance of ACPA^-ve^ RA donors in that study. However, our previous observation of circulating B cells in newly diagnosed, untreated RA patients clearly showed a significant increase in DN2 B cells, which suggests that even at this early stage, DN2 B cells are a prominent B cell subset and may also be expected to be enriched in the synovium (29).

A real strength of this study lies in the opportunity to understand the inter-relatedness of RA synovial B cells. RNA velocity and trajectory inference identified how the clusters of B cells exist in a dynamic continuum. We observed 3 separate lineages, beginning with naïve B cells, cell cycling B cells or the early activation cluster. B cells developed into memory cells before maturing into DN2 B cells and eventually pathogenic ASCs. Clearly the RA synovium can support the generation of ASCs from all three starting points; but all lineages can lead to DN2 B cells.

Previous reports allude to the heterogeneity of autoantibodies that form immune complexes within the joint (8). Similar immune complexes can be found in the joints of patients with osteoarthritis but exist at much higher frequencies in RA synovium (8). The paired BCR sequencing data has strengthened the findings of the RNA velocity analysis and provided an extra layer of detail by tracking B cell clonal lineages based on CDR3 sequence identity. Furthermore, having clonal families containing diverse combinations of B cell subsets including both naïve B cells and ASCs suggests that the immunological response is local to the joint and naïve cells are migrating to the joint where they then become activated. This is supported by another single cell sequencing study of rheumatoid synovial B cells where over 50% of the plasma cells were derived from clonal families that also contained memory B cells (44).

DN2 cells express CD11c, that is driven by the expression of the transcription factor T-bet. The expression of high levels of T-bet in B cells requires the synergistic engagement of the BCR, TLR7 and IFNγ (45). Our data shows that, as well as differentially expressing *ITGAX*, synovial DN2 cells highly express *DUSP4*, that encodes a member of the dual specificity protein phosphatase subfamily. DUSP4 is related to the TLR7/8 cascade thorough its involvement in the MyD88 pathway, further strengthening the link between TLR7 signalling and DN2 B cells (46, 47). Interestingly autophagy plays an essential role in delivering RNA ligands to the endosomes where TLR7 resides (48). Hydroxychloroquine (HCQ), an established therapy for RA, inhibits both TLR7 signalling and autophagy, so HCQ may preferentially constrain the function of DN2 B cells at multiple levels (49, 50). The most differentially expressed gene was *SOX5* gene, that is a member of the Sox family of transcription factors linked to cell fate (51, 52). It has previously been reported that CD11c^+ve^ B cells express both *DUSP4* and *SOX5* and our study supports that observation (53). This transcription factor decreases the proliferative capacity of B cells whilst permitting plasmablast generation (32). Hence expression of SOX5 may licence the evolution of DN2 cells into ASCs.

Synovial DN2 cells can be further divided into two further clusters, that we refer to as DN2-A cells and DN2-B. DN2-A cells are specialised for antigen processing and presentation (APCs), where they likely solicit the appropriate help from synovial T cells. Following this, they downregulate APC function and upregulate expression of *JCHAIN, IGHG1, IGHA1*, and *IL2RG* as they evolve into DN2-B cells, that in turn merges into the ASC cluster. As DN2-A and DN2-B cells function as either APCs or precursors of ASCs, they fulfil at least two of the known criteria for being pathogenic. In contrast, they express few transcripts for cytokines, though we saw no distinct pattern for cytokine production in any of the synovial B cell clusters (Supplementary Fig. S5).

In this study we have shown that DN2 cells are a major precursor to pathogenic ASCs in the RA synovium. We also found that they are primed to present antigen and activate autoreactive T cells. These results fill a gap in our understanding of B cell development in the synovial tissue of RA patients. To fully define the pathogenicity of this B cell subset the antigen specificity of these cells will need to be determined in future studies. Identifying the critical role of DN2 cells in the generation of ASCs provides a novel target for developing future therapeutics.

## Materials and Methods

### Patients

The use of human samples was approved by the South East Scotland Bioresource NHS Ethical Review Board (Ref. 15/ES/0094) and the use of healthy donor blood was approved by AMREC (Ref. 21-EMREC-041). Informed consent was obtained from all study participants prior to sample collection. The patient characteristics are described in Supplementary Table 1.

### Cell Isolation

Peripheral blood mononuclear cells (PBMCs) were isolated from total blood using Histopaque-1077 separation according to the manufacturer’s protocol (Sigma-Aldrich). The PBMCs were collected, washed in PBS, and counted. PBMCs were resuspended in resuspension media (RPMI-1640 (Gibco) supplemented with 2 mM l-glutamine, 100 U ml^-1^ penicillin, and 100 μg ml^-1^ streptomycin, with 40% foetal calf serum (FCS)) then an equal volume of freezing media (resuspension media supplemented with 30% dimethyl sulfoxide (DMSO, Fisher Reagents)) was added to give final density of 10×10^6^/ml in 15% DMSO. The sample was then frozen to −80°C in a Mr Frosty Freezing Container (ThermoFisher Scientific) then stored in liquid nitrogen until further use.

Synovial tissue was collected from RA patients undergoing arthroplasty. The fresh tissue was dissected followed by digestion for 2 hours at 37°C in 1 mg ml^−1^ Collagenase 1 (Sigma-Aldrich). Debris was removed and cells isolated by sequentially passing through 100, 70 and 40μm cell strainers (Corning). The isolated cells were then frozen and stored in the same way as the PBMCs.

### Full spectrum flow cytometry

PBMC and synovial tissue samples were thawed and resuspended according to the 10X protocol (10X Genomics). Briefly, cryovials were removed from liquid nitrogen and thawed rapidly in a water bath at 37°C. The cells were sequentially diluted 5 times with warm media with 1:1 incremental volume addition added in a dropwise manner, gently swirling between each volume. The cell suspension was centrifuged at 230xg for 6 minutes then washed in PBS. A 40μm cell strainer was used to remove debris before the cells were washed and stained in FACS buffer (PBS supplemented with 1% FCS) at room temperature for 20 minutes with fluorescently conjugated antibodies as described in Supplementary Table 2. Cells were washed and re-suspended in FACS buffer and analysed using a Cytek Aurora (Cytek Biosciences) and FCS Express (v7.14, De Novo Software).

### 10X Run and Sequencing

Synovial tissue samples were thawed and resuspended according to the 10X protocol as described above. Cells were washed and then stained in FACS buffer at 4°C for 20 minutes with fluorescently conjugated antibodies as described in Supplementary Table 3. Single CD19^+ve^ DAPI^-ve^ CD3^-ve^ CD14^-ve^ B cells were then sorted with the FACS Aria II and FACSDiva 8.0.1 (BD Biosciences). The full gating strategy is shown in Supplementary Figure S1A. Immediately after sorting the CD19^+ve^ B cells were processed through the Chromium Single Cell Platform using the Chromium Next GEM Single Cell 5’ Kit v2 (10X Genomics, PN-1000265), the Chromium Single Cell Human BCR Amplification Kit (10X Genomics, PN-1000253), and the Chromium Next GEM Chip K Single Cell Kit (10X Genomics, PN-1000287) as per manufacture’s protocol. GEX and V(D)J libraries were then sequenced using the NextSeq2000 at the Wellcome Trust Clinical Research Facility’s Genetics Core in Edinburgh.

### Data pre-processing

The FASTQ files were processed using Cell Ranger (10X Genomics, v7.0.0) and aligned to the GRCh38 reference genome using the cellranger multi command. The raw feature matrices were loaded into R (v4.1.0) using the Seurat (v4.1.0) package (54). Each sample was pre-processed separately before integration using Harmony (v0.1) (55). Cells that expressed fewer than 200 distinct genes, more than 2500 distinct genes, or had more than 5% mitochondrial reads were removed in order to exclude potential multiplets and dead cells. Contaminating non-B cells were then excluded by removing cells expressing CD3E, CD14, GNLY, FCER1A, GCGR3A (CD16), LYZ, or PPBP. Following quality control the three samples had 9,044, 11,840, and 6,169 cells. Gene counts were then log-normalised with default parameters and the top 2,000 variable features for each sample were identified using Seurat’s FindVariableFeatures function. To prevent expression of genes associated with individual BCR clonotypes from influencing dimensionality reduction and clustering, immunoglobulin related genes, including all heavy and light VDJ genes and constant region genes, were removed from the variable features list (1761 variable features remained).

### Clustering, Dimensionality Reduction, and Differential Expression

Unsupervised clustering based on the first 20 principal components of the most variably expressed genes was performed using Seurat::FindNeighbours and Seurat::FindClusters with the resolution 0.5. Clusters were visualised using the manifold approximation and projection (UMAP) method and annotated by analysing the expression of canonical cell markers, SingleR (56), and the use of pre-made gene sets from bulk RNA-seq of sorted B cell populations downloaded from MSigDB (v7.5.1) (57). Differential gene expression between clusters was examined using the MAST test using Seurat::FindMarkers, the results are reported as volcano plots created with EnhancedVolcano (v1.1).

### Module Scores and Gene Set Enrichment Analysis

Module scores were created using the Seurat::AddModuleScore function. The Human SLE DN2 gene signature was generated using the normalised gene expression data from GSE92387 (15), the data was downloaded and analysed using the SARtools R package (58). The top DEGs with >1.5 log2FC were then used as the input list for Seurat::AddModuleScore creating a module score for each cell. The mouse CD21lo CD23lo gene signature was generated using the differentially expressed genes in CD21lo CD23lo cells relative to follicular B cells (full gene list provided in personal correspondence) (19). The module score was created as before but with the top DEGs with >2.5 log2FC. The module scores were then used as features in Seurat::FeaturePlot, and contour plots layered over to show the cells within the top 10% for each module score.

Gene set enrichment analysis (GSEA) was carried out using fgsea (v1.21) (59). Gene Ontology (GO) gene sets were downloaded from MSigDB (v7.5.1). The differentially expressed genes identified by Seurat::FindMarkers were ranked by log2 fold change and used as the input. The GSEA results were reported with the normalised enrichment score (NES).

### Trajectory Inference and RNA Velocity

Trajectory inference was performed using Slingshot (v2.2) (34). Slingshot uses the clusters identified by Seurat in an unsupervised manner to construct a minimum spanning tree which is then converted into smooth lineages aligning cells to a pseudotime trajectory. Trajectory-based differential gene expression analysis was carried out using tradeseq (v1.17) (60) and the lineages identified by Slingshot given as input.

RNA Velocity analysis was performed using Velocyto (v0.17) (61) and scVelo (v 0.2.4) (33). Velocyto was used to identify spliced and unspliced transcripts within the Cellranger output. The resulting loom output file was then merged with the cluster annotation and dimensionality reduction metadata, previously generated in the Seurat analysis, using scVelo. RNA velocities were estimated using the dynamical model from scVelo and projected onto the UMAP.

### BCR Analysis

BCR repertoire data were processed using the Immcantation framework. Initial VDJ assignment was carried out using IgBLAST (v1.18.0) (62) with the aid of the AssignGenes.py wrapper and output was formatted using MakeDb.py, both scripts from Change-O (63). Clustering into clonal groups was also carried out using Change-O. Clones were defined as having identical VJ gene usage and junction length with a normalised Hamming distance threshold of 0.2, as determined by the distToNearest function in the SHazaM R package, used to further partition sequences. Germline sequences for each clonal group were reconstructed using CreateGermlines.py from Change-O. The R package Alakazam was used for clonal lineage reconstruction, lineage topology and repertoire diversity analysis. Mutation counts and frequencies were calculated using a custom Python script, mutation frequency was defined as number of mutations divided by length of sequence. The BCR data, including the mutational loads and clonal assignments, were then integrated with the processed gene expression data in Seurat. Input for the network graphs was generated with custom Python scripts and visualised using Gephi (v0.9.7).

### Statistics

Data were assessed for normality using the Shapiro-Wilk normality test before the appropriate parametric or non-parametric test was chosen. All paired and unpaired Student’s T-tests, Mann-Whitney tests, and Spearman correlations for the full-spectrum flow cytometry data were performed using Prism 9.4 (GraphPad Software Inc.). Use of the ± following mean values indicates the 95% confidence interval.

## Acknowledgements

This work was supported by grants from the Medical Research Council (MR/N013166/1) to EW, the Wellcome Trust (220096/Z/20/Z) to CS, and CSO (TCS/22/03) to MG. The authors thank the QMRI Flow Cytometry and cell sorting facility and the Flow Cytometry Facility at the Institute of Genetics and Cancer for help with the cell sorting. Single-cell sequencing was carried out by the Genetics Core sequencing facility at the Edinburgh Clinical Research Facility, University of Edinburgh. This work has made use of the resources provided by the Edinburgh Compute and Data Facility (ECDF) (http://www.ecdf.ed.ac.uk/).

## Data and Code Availability

Data will be made available at ArrayExpress: upon publication.

All code used for this paper can be found at https://github.com/ElinorWing/DN2_paper.

## Notes

### Competing Interest Statement

The authors have declared no competing interest.

### Summary of Updates

Figure 1 and text

