## Supplementary Fig S1 for "A comprehensive analysis of rheumatoid arthritis B cells reveals the importance of CD11c^+ve^ double-negative-2 B cells as the major synovial plasma cell precursor"

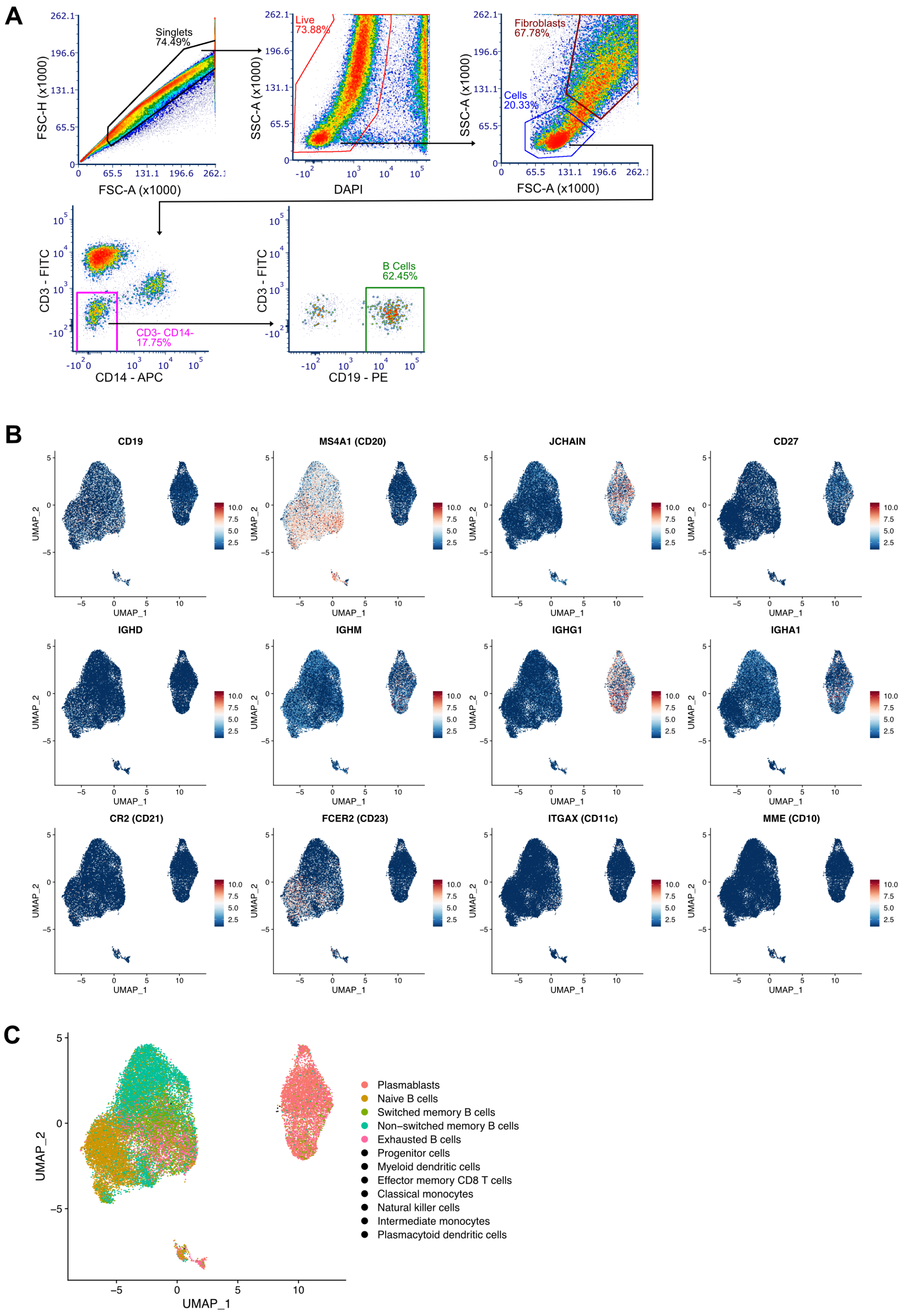


**Supplementary Fig S1 - Isolation and annotation of CD19$^{+ve}$ synovial B cells.**
A) Flow cytometry gating strategy for FACS sorting CD19$^{+ve}$ B cells from human synovial tissue; representative plot from three samples.
B) Selected expression of important genes used for identifying B cell subsets.
C) Annotation from SingleR Monaco Immune Data generated from adult peripheral blood B cell signatures.
