## Supplementary Fig S2 for "A comprehensive analysis of rheumatoid arthritis B cells reveals the importance of CD11c^+ve^ double-negative-2 B cells as the major synovial plasma cell precursor"

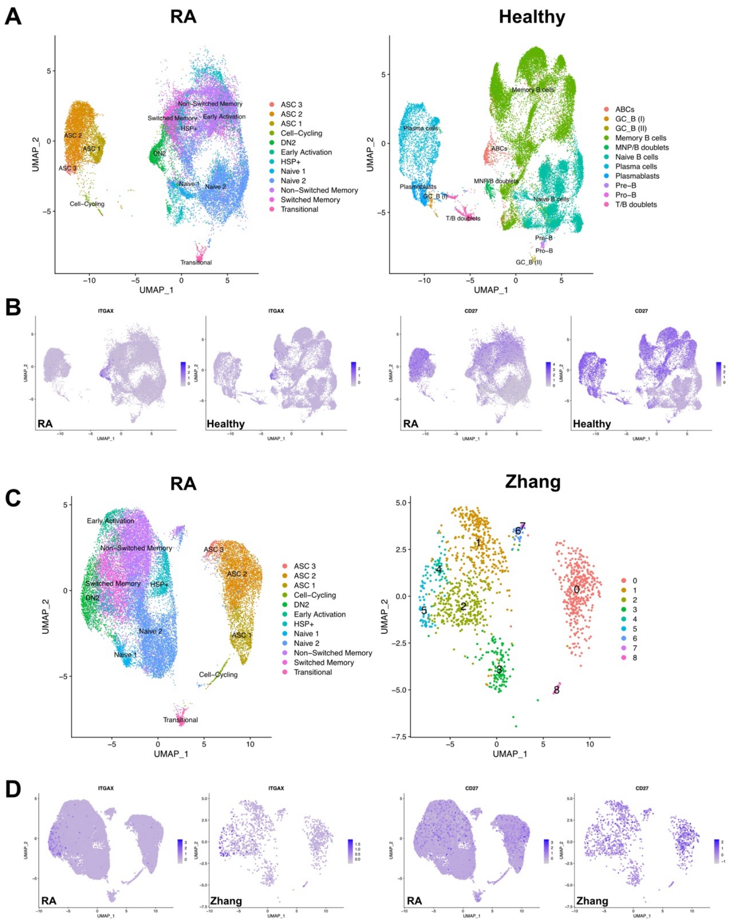


**Supplementary Fig S2: Comparison with publicly available datasets.**
A) Cells from a healthy scRNA-seq data set, including cells from the liver, spleen, and bone marrow were projected onto the same dimensionality-reduced space using UMAP. The cluster labelled as ABCs in this publicly available dataset matches up with our DN2 population.
B) Expression of ITGAX (CD11c) and CD27 in both datasets.
C) Cells from a previously published scRNA-seq dataset from B cells in the synovial tissue of RA patients were projected onto the same dimensionality-reduced space using UMAP. Cluster 5 in this publicly available dataset matches up with our DN2 population and is where the concentration of ITGAX expression is found.
D) Expression of ITGAX (CD11c) and CD27 in both datasets.
