## Supplementary Fig S3 for "A comprehensive analysis of rheumatoid arthritis B cells reveals the importance of CD11c^+ve^ double-negative-2 B cells as the major synovial plasma cell precursor"

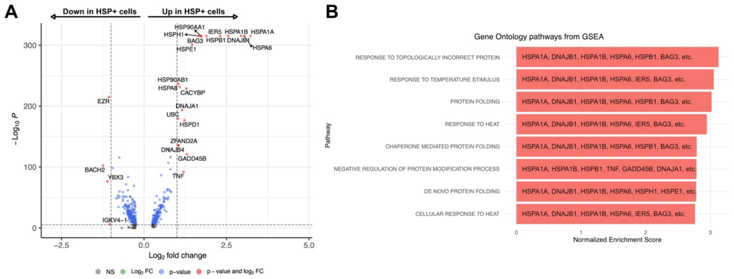


**Supplementary Fig S3: Heat shock protein cluster**

A) Differential expression analysis of HSP+ cells compared to all other non-ASCs.

B) Gene set enrichment analysis using GO Biological Pathways gene sets.
