## Supplementary Fig S4 for "A comprehensive analysis of rheumatoid arthritis B cells reveals the importance of CD11c^+ve^ double-negative-2 B cells as the major synovial plasma cell precursor"

**A**

**B**

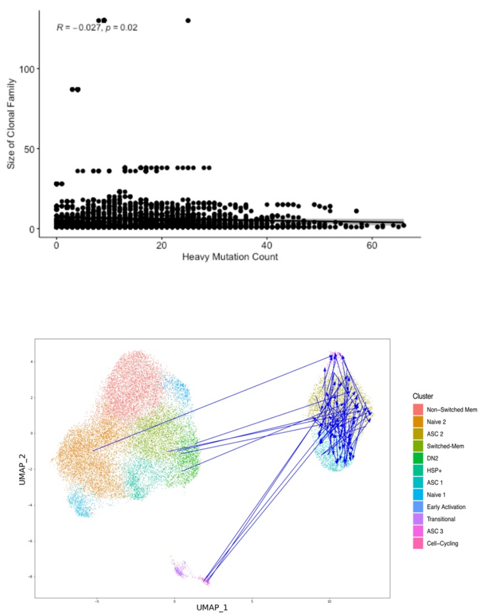


**Supplementary Fig S4: BCR**

A) There was no relationship between the size of the clonal family and the rate of mutation.
B) UMAP with direct relationships ending at an ASC where the direct link indicates an identical heavy CDR3 amino acid sequence between the cells (Fig. 6C and D). These direct links, ending at an ASC, originated from 4 DN2 cells, that in turn emerged from 2 Switched Memory cells, 1 Naïve-2 cell and 4 Cell-Cycling cells, demonstrating that DN2 cells are the primary precursor to ASCs.
