## Supplementary Fig S5 for "A comprehensive analysis of rheumatoid arthritis B cells reveals the importance of CD11c^+ve^ double-negative-2 B cells as the major synovial plasma cell precursor"

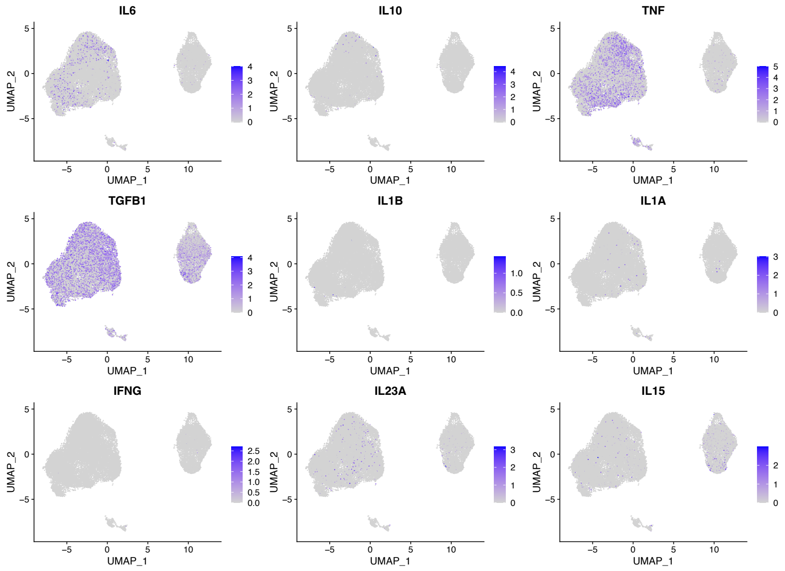


**Supplementary Fig S5: Cytokine gene expression patterns.**
Expression of important cytokine genes in rheumatoid arthritis.
