## Supplementary Table 1 for "A comprehensive analysis of rheumatoid arthritis B cells reveals the importance of CD11c^+ve^ double-negative-2 B cells as the major synovial plasma cell precursor"

Supplementary Table 1 – Characteristics of patients used for full spectrum flow cytometry and the scRNA-seq.

| | RA Patients (n=29) | Paired Synovium and Blood (n=5) | Arthroplasty Patients (n=3) |
|---|---|---|---|
| Age (year, IQR) | 57.5 (51.5-65) | 58.9 (49.8-68.6) | 66.8 (66.6-68.1) |
| Female (%) | 21 (72.4) | 4 (80) | 3 (100) |
| CCP (IU/ml, IQR) | 309 (120-340) *[1] | 219 (105.5-323) *[4] | 473 (336.6-536.5) |
| RF (IU/ml, IQR) | 100 (37.5-205) *[2] | 314.72 (200-500) | 326.5 (263.3-389.75) *[5] |
| Baseline DAS28 (IQR) | 5.16 (2.95-7.36) *[3] | N/A | N/A |
| DMARD treatment | | | |
| -Methotrexate | 7 | 2 | 1 |
| -Hydroxychloroquine | 5 | 1 | 0 |
| -Sulfasalazine | 2 | 2 | 0 |
| -Leflunomide | 1 | 0 | 0 |
| Number of concurrent DMARDS | | | |
| -One | 8 | 1 | 1 |
| -Two | 2 | 2 | 0 |
| -Three | 1 | 0 | 0 |
| Biologic treatment | | | |
| -Etanercept | 0 | 1 | 0 |

*[1] n = 22, *[2] n = 21, *[3] n = 27, *[4] n = 3, *[5] n = 2.
