## Supplementary Table 2 for "A comprehensive analysis of rheumatoid arthritis B cells reveals the importance of CD11c^+ve^ double-negative-2 B cells as the major synovial plasma cell precursor"

Supplementary Table 2 - Antibody reagents for FACS sorting.

| Antibody (Clone) | Fluorochrome | Isotype | Source | Dilution |
|---|---|---|---|---|
| Anti-human CD19 (HIB19) | PE | Mouse IgG1, κ | BioLegend | 1:50 |
| Anti-human CD3 (HIT3a) | FITC | Mouse IgG2a, κ | BioLegend | 1:50 |
| Anti-human CD14 (63D3) | APC | Mouse IgG1, κ | BioLegend | 1:50 |
| DAPI | | | Sigma-Aldrich | 1:10000 |
