## Supplementary Table 3 for "A comprehensive analysis of rheumatoid arthritis B cells reveals the importance of CD11c^+ve^ double-negative-2 B cells as the major synovial plasma cell precursor"

Supplementary Table 3 - Antibodies used in full spectrum flow cytometry staining.

| Antibody (Clone) | Fluorochrome | Isotype | Source | Panel | Dilution |
|---|---|---|---|---|---|
| Anti-human CD10 (HI10a) | Brilliant Violet 480 | Mouse IgG1, κ | BD Biosciences | 1 & 2 | 1:67 |
| Anti-human CD11c (3.9) | Brilliant Violet 785 | Mouse IgG1, κ | BioLegend | 1 & 2 | 1:50 |
| Anti-human CD138 (1D4) | PE-Cy5 | Mouse IgG1 | Stratech | 1 | 1:50 |
| Anti-human CD138 (MI15) | PE/Cy5 | Mouse IgG1, κ | AAT Bioquest | 2 | 1:50 |
| Anti-human CD14 (63D3) | BV510 | Mouse IgG2a, κ | BioLegend | 1 | 1:50 |
| Anti-human CD184 (CXCR4) (12G5) | PerCP-eFluor™ 710 | Mouse IgG2a, κ | ThermoFisher Scientific | 2 | 1:50 |
| Anti-human CD185 (CXCR5) (MU5UBEE) | Super Bright 436 | Mouse IgG2b, κ | ThermoFisher Scientific | 1 & 2 | 1:50 |
| Anti-human CD19 (HIB19) | Brilliant Violet 711 | Mouse IgG1, κ | BioLegend | 2 | 1:50 |
| Anti-human CD19 (HIB19) | PE/Dazzle™ 594 | Mouse IgG1, κ | BioLegend | 1 | 1:50 |
| Anti-human CD20 (2H7) | eFluor 450 | Mouse IgG2b, κ | ThermoFisher Scientific | 1 & 2 | 1:50 |
| Anti-human CD21 (Bu32) | PE/Cyanine7 | Mouse IgG1, κ | BioLegend | 1 & 2 | 1:50 |
| Anti-human CD23 (M-L233) | Brilliant Violet 750 | Mouse IgG1, κ | BD Biosciences | 1 & 2 | 1:67 |
| Anti-human CD24 (ML5) | Brilliant Violet 650 | Mouse IgG2a, κ | BD Biosciences | 1 & 2 | 1:50 |
| Anti-human CD27 (M-T271) | APC | Mouse IgG1, κ | BD Biosciences | 2 | 1:20 |
| Anti-human CD3 (UCHT1) | Alexa Fluor 532 | Mouse IgG1, κ | ThermoFisher Scientific | 1 & 2 | 1:50 |
| Anti-human CD307e (FcRL5) (509f6) | PE | Mouse IgG2a, κ | BioLegend | 1 & 2 | 1:50 |
| Anti-human CD38 (HIT2) | Brilliant Violet 605 | Mouse IgG1, κ | BioLegend | 1 & 2 | 1:50 |
| Anti-human CD39 (A1) | PE/Fire™ 810 | Mouse IgG1, κ | BioLegend | 2 | 1:50 |
| Anti-human CD40 (5C3) | Alexa Fluor 700 | Mouse IgG1, κ | BioLegend | 2 | 1:50 |
| Anti-human CD45RB (MT4) | BV711 | Mouse IgG1, κ | BD | 1 | 1:100 |
| Anti-human CD5 (L17F12) | PE/Dazzle™ 594 | Mouse IgG2a, κ | BioLegend | 2 | 1:50 |
| Anti-human CD73 (Ecto-5'-nucleotidase) (AD2) | FITC | Mouse IgG1, κ | BioLegend | 1 & 2 | 1:50 |
| Anti-human CD86 (BU63) | Brilliant Violet 421 | Mouse IgG1, κ | BioLegend | 1 & 2 | 1:50 |
| Anti-human CD95 (Fas) (DX2) | PE/Fire™ 640 | Mouse IgG1, κ | BioLegend | 2 | 1:50 |
| Anti-human CD95 (Fas) (DX2) | AF700 | Mouse IgG1, κ | BioLegend | 1 | 1:50 |
| Anti-human HLA-DR (L243) | APC/Fire™ 810 | Mouse IgG2a, κ | BioLegend | 1 & 2 | 1:50 |
| Anti-human IgA (IS11-8E10) | VioGreen | Mouse IgG1, κ | Miltenyi Biotec | 2 | 1:100 |
| Anti-human IgD (IA6-2) | APC/Fire™ 750 | Mouse IgG2a, κ | BioLegend | 2 | 1:50 |
| Anti-human IgD (IA6-2) | APC-Cy7 | Mouse IgG2a, κ | BioLegend | 1 | 1:50 |
| Anti-human IgG (M1310G05) | Alexa Fluor 647 | Rat IgG2a, κ | BioLegend | 1 & 2 | 1:50 |
| Anti-human IgM (MHM-88) | Brilliant Violet 570 | Mouse IgG1, κ | BioLegend | 2 | 1:50 |
| Anti-human IgM (SA-DA4) | PerCP-eF710 | Mouse IgG1, κ | ThermoFisher | 1 | 1:50 |
| Fixable Viability Kit | Zombie NIR™ | | BioLegend | 1 & 2 | 1:1000 |
