## Supplementary Table 4 for "A comprehensive analysis of rheumatoid arthritis B cells reveals the importance of CD11c^+ve^ double-negative-2 B cells as the major synovial plasma cell precursor"

Supplementary Table 4 - Subset proportions for all known B cell subsets in the blood. Mann-Whitney test for non-normally distributed, unpaired data. t test for normally distributed, unpaired data.

| B Cell Population | | Markers | Healthy (%, IQR) n=15 | RA (%, IQR) n=29 | P value |
|---|---|---|---|---|---|
| Naive | | $IgD^{+ve}$ $CD27^{-ve}$ | 70.3 (14.4) | 72.3 (8.2) | 0.5837 |
| Unswitched Memory | | $IgD^{+ve}$ $CD27^{+ve}$ | 12.3 (8.83) | 10.3 (5.54) | 0.4188 |
| Switched Memory | | $IgD^{-ve}$ $CD27^{+ve}$ | 13.4 (6.9) | 13.2 (6.9) | 0.9233 |
| Double Negative | | $IgD^{-ve}$ $CD27^{-ve}$ | 11.4 (5.25) | 13.5 (5.0) | 0.2721 |
| Transitional | T1 | $IgD^{+ve}$ $CD27^{-ve}$ $CD38^{high}$ $CD24^{high}$ | 3.68 (2.04) | 3.58 (1.9) | 0.8871 |
| | T2-MZP | $IgD^{+ve}$ $CD27^{-ve}$ $CD38^{high}$ $CD24^{high}$ $CD21^{high}$ | 0.227 (0.27) | 0.226 (0.27) | 0.3934 |
| | T3 | $IgD^{+ve}$ $CD27^{-ve}$ $CD38^{+ve}$ $CD24^{+ve}$ $CD21^{+ve}$ | 1.75 (1.95) | 1.67 (1.35) | 0.6727 |
| Naive | Resting | $CD38^{+ve}$ $CD24^{+ve}$ $CD21^{+ve}$ | 48.6 (13.5) | 47.1 (13.1) | 0.7772 |
| | Activated | $CD38^{-ve}$ $CD24^{-ve}$ $CD21^{-ve}$ | 1.13 (0.889) | 2.35 (1.52) | **0.0091** |
| | Anergic | $CD38^{+ve/low}$ $CD24^{+ve}$ $CD21^{-ve}$ | 6.61 (4.0) | 5.79 (2.2) | 0.6773 |
| Memory | Unswitched | $CD38^{+ve/low}$ $CD24^{+ve}$ $CD21^{+ve}$ | 10.1 (8.2) | 8.09 (4.6) | 0.3139 |
| | $IgM^{+ve}$ Only | $IgD^{-ve}$ $IgM^{+ve}$ $CD27^{+ve}$ $CD38^{+ve/low}$ $CD24^{+ve}$ $CD21^{+ve}$ | 2.86 (2.1) | 1.32 (0.718) | **0.0008** |
| | Switched Resting | $CD38^{+ve/low}$ $CD24^{+ve}$ $CD21^{+ve}$ | 6.92 (4.92) | 5.36 (3.5) | 0.2855 |
| | Switched Active | $CD38^{-ve}$ $CD24^{-ve}$ $CD21^{-ve}$ | 0.607 (0.50) | 1.24 (1.07) | 0.0943 |
| Double Negative | DN1 | $CD38^{+ve}$ $CD24^{+ve}$ $CD21^{+ve}$ | 7.28 (3.9) | 6.78 (3.94) | 0.3479 |
| | DN2 | $CD38^{-ve}$ $CD24^{-ve}$ $CD21^{-ve}$ | 0.757 (0.77) | 1.51 (0.59) | **0.0003** |
| ASC | | $CD27^{high}$ $CD20^{-ve}$ | 1.33 (0.92) | 2.43 (2.0) | 0.5278 |
| B1 | | $IgD^{high}$ $CD27^{+ve}$ | 0.691 (0.92) | 0.933 (0.81) | 0.6027 |
